# *EARLY MATURITY 7* modifies the circadian clock and photoperiod sensitivity in barley

**DOI:** 10.1101/2023.06.22.546112

**Authors:** Gesa Helmsorig, Agatha Walla, Thea Rütjes, Gabriele Buchmann, Rebekka Schüller, Götz Hensel, Maria von Korff

**Author notes:** Corresponding author: Maria von Korff.

## Abstract

Breeding for variation in photoperiod response is crucial to adapt crop plants to novel environments. Plants measure changes in day length by the circadian clock, an endogenous timekeeper that allows plants to anticipate changes in diurnal and seasonal light-dark cycles. Here, we describe the *early maturity 7* (*eam7*) mutation in barley, which interacts with natural variation at *PHOTOPERIOD 1* (*Ppd-H1*) to cause early flowering independent of the photoperiod. We identify *LIGHT-REGULATED WD 1 (LWD1)* as a putative candidate to underly the *eam7* locus in barley as supported by genetic mapping and CRISPR-Cas9 generated *lwd1* mutants. Mutations in *eam7* cause a significant phase advance and a misregulation of core clock and clock output genes under diurnal conditions. Early flowering correlated with an upregulation of *Ppd-H1* during the night and consequent induction of the florigen *FLOWERING LOCUS T1* under short days. We propose that *EAM7* controls photoperiodic flowering in barley by controlling the light input into the clock and diurnal expression patterns of the major photoperiod response gene *Ppd-H1*.

## Introduction

Flowering time significantly impacts crop yield and is thus a vital breeding target to produce novel varieties better adapted to diverse and changing climatic conditions (Cockram et al., 2007). Climate models predict an increase in global temperatures and extreme weather events. Breeding for early maturing varieties is among the most effective strategies to improve adaptation to short growing seasons with terminal stress such as heat and drought events (Tewolde et al., 2006; He et al., 2022). Identification and characterization of novel alleles conferring early maturity greatly support these efforts.

The temperate crop barley (*Hordeum vulgare*) is among the most widely grown cereals with superior adaptation to marginal stress-prone agricultural lands (von Korff et al., 2008). Like most temperate cereals, barley is a quantitative long-day (LD) species that accelerates reproductive development with increasing photoperiods. In contrast, short photoperiods delay or even impair floral development (Digel et al., 2015). *PHOTOPERIOD 1* (*Ppd-H1*), orthologous to *PSEUDO-RESPONSE-REGULATOR (PRR)* genes in *Arabidopsis thaliana*, has been identified as the central gene in photoperiodic flowering in barley (Laurie et al., 1995). Natural variation at *Ppd-H1* determines variation in reproductive development under LDs. The wild-type *Ppd-H1* allele, prevalent in wild and winter barley, induces rapid floral development in response to long days (Turner et al., 2005; Jones et al., 2008). A single amino acid change in the conserved CCT domain of *ppd-H1* delays flowering. This allele was selected and is prevalent in spring barley varieties in central and northern Europe (Jones et al., 2008). *Ppd-H1* initiates flowering by inducing the expression of *FLOWERING LOCUS T1 (FT1)* in the leaves (Turner et al., 2005). *FT1* is orthologous to florigen *FT* in Arabidopsis and *Hd3a* in rice, which move as proteins from the leaf to the shoot apical meristem and induce the transition from vegetative to reproductive growth (Kardailsky et al., 1999; Corbesier et al., 2007; Tamaki et al., 2007). In barley, increased *FT1* expression in the leaf correlates with early flowering and the upregulation of floral inducers such as *MADS*-box genes *VERNALIZATION 1 (VRN1), BM3,* and *BM8* in the leaf and meristem (Turner et al., 2005; Digel et al., 2015).

Mutations conferring photoperiod-independent early flowering, so-called *early maturity (eam)* loci, have been used in Scandinavian breeding programs since the 1960s to enable a geographic range extension of barley cultivation to areas with short growing seasons (Lundqvist, 2009). Several genes that underly *eam* loci have been identified in the last decade: *eam5* was identified as a gain-of-function mutation in *PHYTOCHROME C (PHYC), eam8* as a knock-out mutation in *EARLY FLOWERING 3 (ELF3)* and *eam10* as a single amino acid exchange in *LUX ARRHYTHMO 1 (LUX1)* (Faure et al., 2012; Zakhrabekova et al., 2012; Campoli et al., 2013; Pankin et al., 2014). *ELF3* and *LUX1* are both components of the evening complex (EC) of the circadian clock. Plants carrying *eam* mutations are characterized by reduced photoperiod sensitivity and accelerated flowering in inductive LD and non-inductive short days (SD).

Differences in photoperiod are perceived by photoreceptors, which transmit this information to the circadian clock. The circadian clock is an endogenous timekeeper that allows organisms to anticipate seasonal and daily changes in light-dark rhythms. The core circadian clock is largely conserved across eudicots and monocots (Song et al., 2010). It comprises three interlocking, negative feedback loops of transcriptional repressors that are expressed sequentially during a 24 h period and repress previous and subsequent clock components. Transcription factors *CIRCADIAN CLOCK ASSOCIATED 1 (CCA1)* and *LATE ELONGATED HYPOCOTYL (LHY*) are expressed first in the morning, followed by *PRR9, 7,* and *5* during the progressing day, and *TOC1* (*PRR1*) at dusk. *CCA1/LHY* repress the expression of *PRR* genes, which, in turn, suppress *CCA1/LHY*. During the night, EC genes *ELF3, EARLY FLOWERING 4 (ELF4),* and *LUX* are expressed, repressing *PRR* genes and *CCA1/LHY* (Farré et al., 2005; Kikis et al., 2005; Huang et al., 2012; Mizuno et al., 2014). In Arabidopsis, the clock controls the diurnal expression pattern of the central photoperiod response gene *CONSTANS* (*CO)*, resulting in *CO* expression peaking at the end of the light period in LDs, but in the dark under SDs (Suárez-López et al., 2001). The coincidence of *CO* expression with the light period is necessary for protein stabilization and the expression of the florigen *FT* under LDs (Suárez-López et al., 2001; Valverde et al., 2004; Sawa et al., 2007; Jang et al., 2008). By contrast, in temperate monocots, the length of the night, rather than the length of the day, is critical for the perception of inductive photoperiods (Pearce et al., 2017; Gao et al., 2019). The length of the night is measured by the phytochromes *PHYB* and *PHYC,* which are necessary for the light activation of *Ppd-H1*, and mutations in either of these genes result in the downregulation of *Ppd-H1* and very late flowering (Chen et al., 2014; Kippes et al., 2020). In addition, *PHYB* and *PHYC* control the light-induced degradation of *ELF3* (Gao et al., 2019; Alvarez et al., 2023). *ELF3*, proposedly together with EC members *LUX1* and *ELF4*, binds to the promoter of *Ppd-H1* during the night to repress its expression (Gao et al., 2019; Andrade et al., 2022; Alvarez et al., 2023). While clock mutants are characterized by loss of circadian transcriptome oscillations and severely perturbed clock functions (Müller et al., 2016; Müller et al., 2020), they have been instrumental in expanding the cultivation of many crops to new geographic regions with altered annual patterns of temperature and photoperiod (McClung, 2021). It is thus interesting to identify novel genes and alleles and decipher the molecular networks important for photoperiod response in crops.

Here, we describe the *early maturity 7 (eam7)* locus, originally identified as a natural mutation in the cultivar Atlas, that causes early flowering under non-inductive SD conditions (Stracke and Börner, 1998). We identify *LIGHT-REGULATED WD 1 (LWD1)* as a putative candidate to underly the *EAM7* locus in barley as supported by genetic mapping and CRISPR-Cas9 generated *lwd1* mutants. We demonstrate that *eam7* interacts with natural variation at *Ppd-H1* to promote flowering under non-inductive photoperiods and affects plant architecture, spike development, spike fertility, and grain set. We propose that *EAM7* controls photoperiodic flowering in barley by repressing *Ppd-H1* at night, possibly mediated through *ELF3*. Mutations in *EAM7* cause a significant phase advance and a downregulation of core clock and clock output genes under diurnal conditions. We thus suggest that *EAM7/LWD1* controls photoperiod response by modifying the light entrainment of the clock and clock gene expression.

## Results

### *eam7* accelerates reproductive development in long-day and short-day conditions

We investigated the effects of *eam7* and *Ppd-H1* on flowering time under long and short photoperiods. For this purpose, the spring cultivar Bowman (BW) and three derived introgression lines were cultivated under controlled conditions under long-day (LD, 16 h light / 8 h dark, 20°C / 16°C) or short-day (SD, 8 h light / 16 h dark, 20°C / 16°C) conditions to score flowering time. BW carries a natural mutation in the CCT domain of *Ppd-H1* that delays flowering time under LD conditions (Turner et al., 2005). The derived introgression line BW*(Ppd- H1)* carries a wild-type *Ppd-H1* allele and is early flowering under LDs (Druka et al., 2011). In addition, we used BW*(eam7),* an introgression line with the *eam7.g* mutation that causes early maturity under LD and SD (Stracke and Börner, 1998; Druka et al., 2011). We crossed BW*(Ppd- H1)* and BW*(eam7)* to generate a line with a wild-type *Ppd-H1* allele and the mutation at *eam7*, which we termed BW*(Ppd-H1,eam7)*.

Under LD, both introgression lines with a wild-type *Ppd-H1* allele, BW*(Ppd-H1)* and BW(*Ppd- H1,eam7*), flowered 27 and 26 days after emergence (DAE), respectively, and therefore significantly earlier than BW and BW*(eam7),* which flowered 41 DAE (Fig. 1A). No significant differences in time to flowering were observed between BW and BW*(eam7)* and between BW*(Ppd-H1)* and BW(*Ppd-H1,eam7*) under LDs. Consequently, *Ppd-H1* but not *eam7* controlled time to flowering under LDs. Under SD, BW*(eam7)* flowered 90 DAE and thus significantly earlier than BW and BW*(Ppd-H1),* which flowered on average 98 and 101 DAE, respectively (Fig. 1B). However, 10% of BW and 22% of BW*(Ppd-H1)* plants had not flowered until the experiment was stopped at 125 days. BW*(Ppd-H1,eam7)* exhibited the fastest development and flowered 38 DAE and thus 52 days earlier than BW*(eam7)*, indicating that *eam7* and *Ppd-H1* interacted to accelerate flowering under SD.

**Figure 1:**
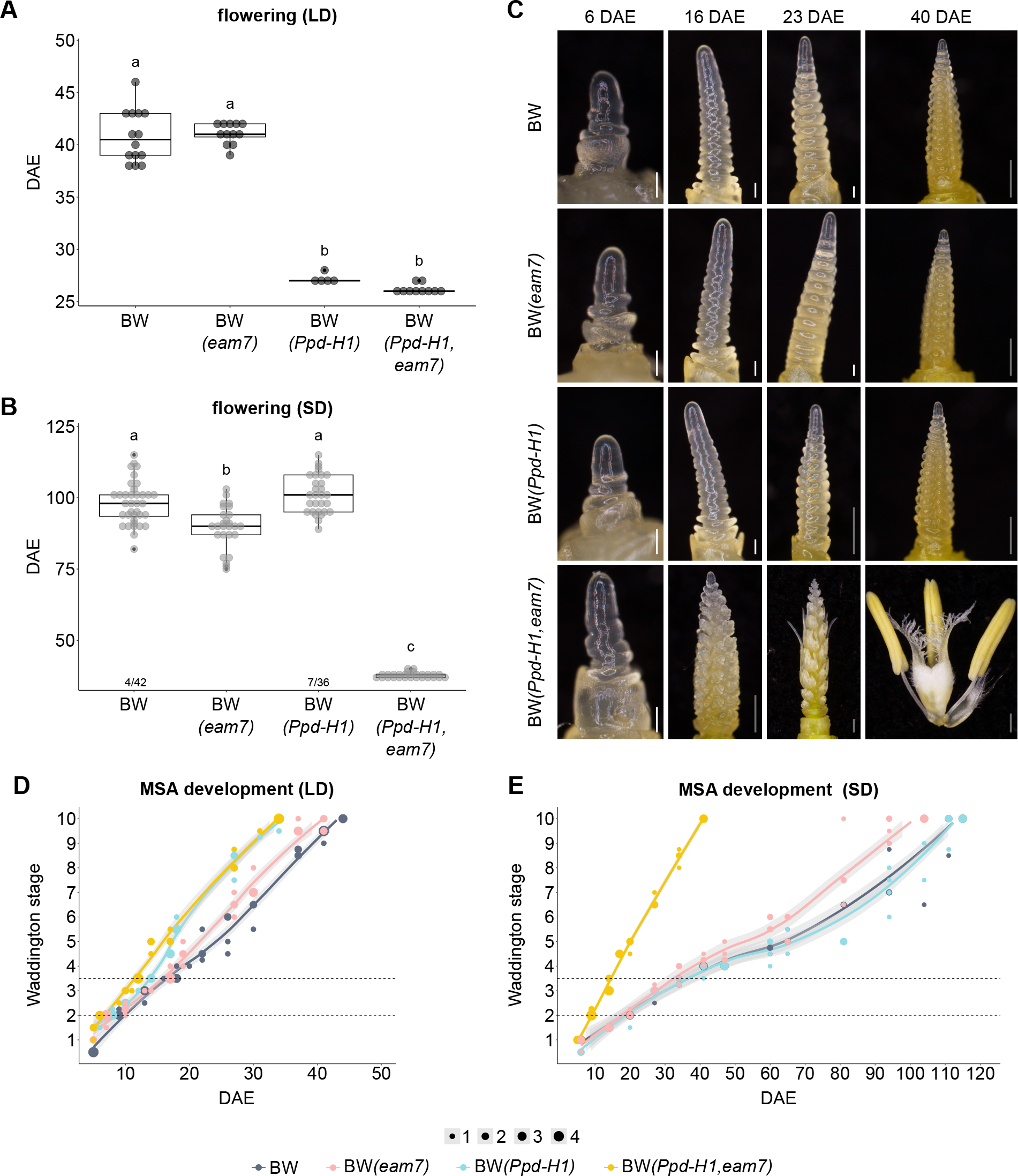
Effects of *eam7* and *Ppd-H1* on plant reproductive development. A, B. Flowering time of spring cultivar Bowman (BW) and introgression lines BW*(eam7)*, BW*(Ppd-H1),* and BW*(Ppd-H1,eam7)* was scored in days after emergence (DAE) under long days **(A)** and short days **(B)**. Significance levels were determined by one-way ANOVA and subsequent Tukey’s test, *p* ≤ 0.05, n = 5-42 plants. Numbers below boxplots in (B) indicate the number of plants that did not flower until the end of the experiment (125 DAE) compared to the total number of plants scored. **C** Inflorescences of BW, BW*(eam7)*, BW*(Ppd-H1),* and BW*(Ppd-H1,eam7)* plants grown under short day. The main culm was dissected 6, 16, 23, and 40 DAE. White scale bars indicate 100 μm, and grey scale bars 500 μm. **D, E** Main shoot apex (MSA) development under long-day (**D)** and short-day **(E)** conditions according to the scale by Waddington et al. (1983) by DAE. Dot sizes indicate the number of plants per data point (1-4), and grey areas show a 95% confidence interval of a polynomial regression (Loess smooth line). Horizontal lines indicate the start of spikelet initiation (W2.0) and the start of floral development (W3.5).

We then analyzed which stages of reproductive development were affected by variation at *EAM7* and *Ppd-H1.* The main shoot apices (MSA) of plants grown under LD and SD were dissected over development and scored according to the scale by Waddington et al. (1983). The Waddington scale rates the development based on the carpel of the most advanced floret of the spike. This way, MSA development can be categorized into vegetative growth (W1.0-W2.0), in which leaf primordia are initiated, early reproductive growth (W2.0-W3.5) with the initiation of spikelet meristems (SM), and late reproductive growth and floral development until anthesis and pollination (W3.5-W10.0).

Under LD, BW*(Ppd-H1,eam7)* transitioned to reproductive growth (W2.0) six DAE and thus two days earlier than BW*(Ppd-H1)* and BW*(eam7)* and three days earlier than BW (Fig. 1D). While BW*(Ppd-H1,eam7)* developed faster than BW*(Ppd-H1)* during early reproductive growth, their development synchronized during floral organ growth, and both genotypes reached pollination (W10.0) 34 DAE. In contrast, *BW(eam7)* and BW developed similarly during early reproductive growth but BW*(eam7)* development accelerated after carpel initiation (W4.5), and plants reached pollination three days earlier than BW(Fig. 1D). Under SDs, the MSA of BW*(Ppd- H1,eam7)* transitioned to reproductive growth nine DAE and thus 11 days earlier compared to the other three genotypes (Fig. 1E). BW*(Ppd-H1,eam7)* plants displayed a linear reproductive development under SD so that pollination (W10.0) occurred 41 DAE and thus only seven days later compared to LDs (Fig. 1C). By contrast, BW, BW*(Ppd-H1)*, and BW*(eam7)* showed a substantial delay in floral development after carpel initiation (W4.5). However, floral development was still faster in BW*(eam7)* compared to BW and BW*(Ppd-H1),* and pollination occurred at 98 DAE, compared to 111 DAE in BW and BW*(Ppd-H1)* (Fig. 1E).

Consequently, under LDs, *eam7* accelerated spikelet initiation in the background of *Ppd-H1* and floral growth in the background of *ppd-H1*. Under SD, *eam7* strongly accelerated all stages of reproductive development in the background of the wild-type *Ppd-H1* allele. In contrast, it only accelerated floral organ growth by a few days in the background of the mutated *ppd-H1* allele.

Next, we investigated the effects of variation at *EAM7* and *Ppd-H1* on inflorescence architecture by scoring the initiation of SM and the number of florets and grains on the main spike. Under LD, BW produced the highest number of florets and grains per main spike, followed by BW*(eam7)*, BW*(Ppd-H1),* and BW*(Ppd-H1,eam7)* (Supplemental Fig. 2A-C, E). This result was correlated to a longer duration of SM initiation and a higher number of total SM initiated in BW (Supplemental Fig. S1A, Supplemental Table S1). Spike fertility, the number of grains per florets on the spike, was close to 75% for BW, BW*(Ppd-H1),* and BW*(eam7)* in contrast to BW*(Ppd- H1,eam7)* with reduced fertility of only 25% (Fig. 2D). The low number of grains per spike in BW*(Ppd-H1,eam7)* was thus caused by a reduced number of initiated SM and reduced floret fertility. Under SD, BW*(Ppd-H1,eam7)* still initiated significantly less SM compared to the other genotypes, spike fertility, however, was relatively higher in BW*(eam7)* and BW*(Ppd-H1,eam7)* compared to BW*(Ppd-H1)* and BW (Fig. 2J), (Supplemental Fig. S1B, Supplemental Table S1).

**Figure 2:**
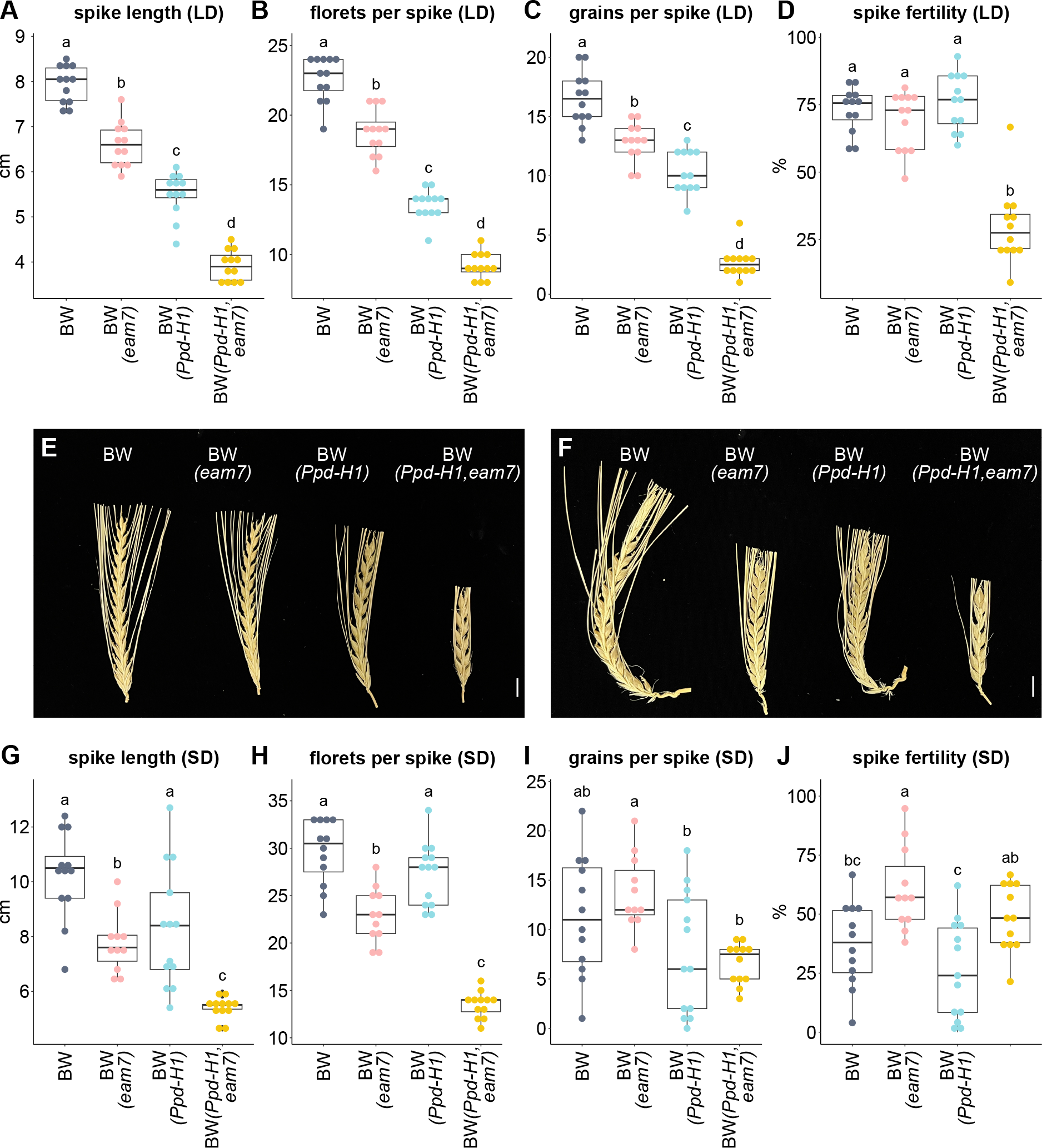
Effect of *eam7* on fertility under LD and SD. Spike length (in cm), floret and grain number on the spike, and spike fertility (in %) were scored on the main culm of plants grown under long days **(A-D)** or short days **(G-J)**. Spike fertility was calculated by dividing the number of grains by the number of florets on the main spike. Significance levels were determined by one-way ANOVA and subsequent Tukey’s test, *p* ≤ 0.05, n = 11-13 plants. **E, F** Representative images of fully developed spikes from the main culm of plants grown under long days **(E)** or short days **(F)**. Scale equals 1 cm.

The effects of *Ppd-H1* and *EAM7* on inflorescence architecture and floret fertility thus differed between photoperiods (Fig. 2F, G-I).

In addition, we determined the effects of *EAM7* on plant height, tiller number, and number of leaves on the main culm under LD and SD. Under LD, variation in shoot architecture was mainly affected by *Ppd-H1*, as BW*(Ppd-H1)* and BW*(Ppd-H1,eam7)* transitioned to reproductive growth earlier and thus produced fewer leaves on the main culm, fewer tillers and grew less tall compared to BW and BW*(eam7)* (Supplemental Fig. S2A-C). Similarly, under SD, the fast-developing BW*(Ppd-H1,eam7)* plants produced significantly fewer leaves and tillers and were characterized by shorter plant height compared to the other three genotypes with a slower reproductive development (Supplemental Fig. S2D-J). We also scored flag leaf size under SD and found that flag leaf length and width were reduced in BW*(Ppd-H1,eam7)* and BW*(eam7)* compared to BW and BW*(Ppd-H1)* (Supplemental Fig. S2K-M).

In conclusion, under LD, *Ppd-H1* had a significant effect on developmental timing, while variation at *EAM7* only had minor effects on floral growth and the timing of pollination. *Ppd-H1* strongly affected the rate and duration of SM initiation and floret and grain number, further modulated by variation at *EAM7.* Under SD conditions, *eam7* in the background of *Ppd-H1* strongly accelerated reproductive development and caused near-day-length neutrality and early flowering independent of the photoperiod. *EAM7* interacted with *Ppd-H1* to control SM initiation and floret fertility under SD. In the background of a mutated *ppd-H1* allele, *eam7* did not affect SM initiation but still affected floret, grain number, and spike fertility. Early flowering decreased plant height, the number of leaves and tillers, and flag leaf size under LDs and SDs.

### The expression pattern of circadian clock genes is altered in *eam7* plants under SD

Day-length neutrality and early flowering under short-day conditions have been associated with genetic variation in circadian clock genes and phytochromes in cereal crops (Faure et al., 2012; Campoli et al., 2013; Pankin et al., 2014; Müller et al., 2020; Alvarez et al., 2023). We therefore tested if the diurnal expression pattern of phytochromes and core clock genes was altered by *eam7*. As the most substantial effects of *eam7* on development were observed under SD, we tested the diurnal expression of phytochromes and core clock genes under SD conditions.

The expression patterns of *PHYB, PHYC*, and clock genes were strongly altered in BW*(eam7)* and BW*(Ppd-H1,eam7)* compared to BW and BW*(Ppd-H1)* (Fig. 3). At the same time, expression patterns of phytochromes and clock genes did not differ between BW and BW*(Ppd-H1)* and between BW*(eam7)* and BW*(Ppd-H1,eam7),* suggesting that *eam7*, but not *Ppd-H1* had a major impact on the diurnal expression patterns. *PHYB/C* and *ELF3* were significantly downregulated at their expression peaks in BW*(eam7)* and BW*(Ppd-H1,eam7)* compared to BW and BW*(Ppd-H1)* (Fig. 3A-C). Furthermore, expression peaks of the evening-expressed clock genes *LUX1, PRR59, GI, PRR1,* and *LHY* were advanced by 2-4 h and strongly reduced in BW*(eam7)* and BW*(Ppd-H1,eam7)* (Fig. 3D, G, I-K). The night-time repression of *Ppd-H1, PRR73, PRR59,* and *PRR95* was released in BW*(eam7)* and BW*(Ppd-H1,eam7),* which resulted in high transcript levels in the night and morning (Fig. 3E-H). This effect was particularly prominent for *Ppd-H1*.

**Figure 3:**
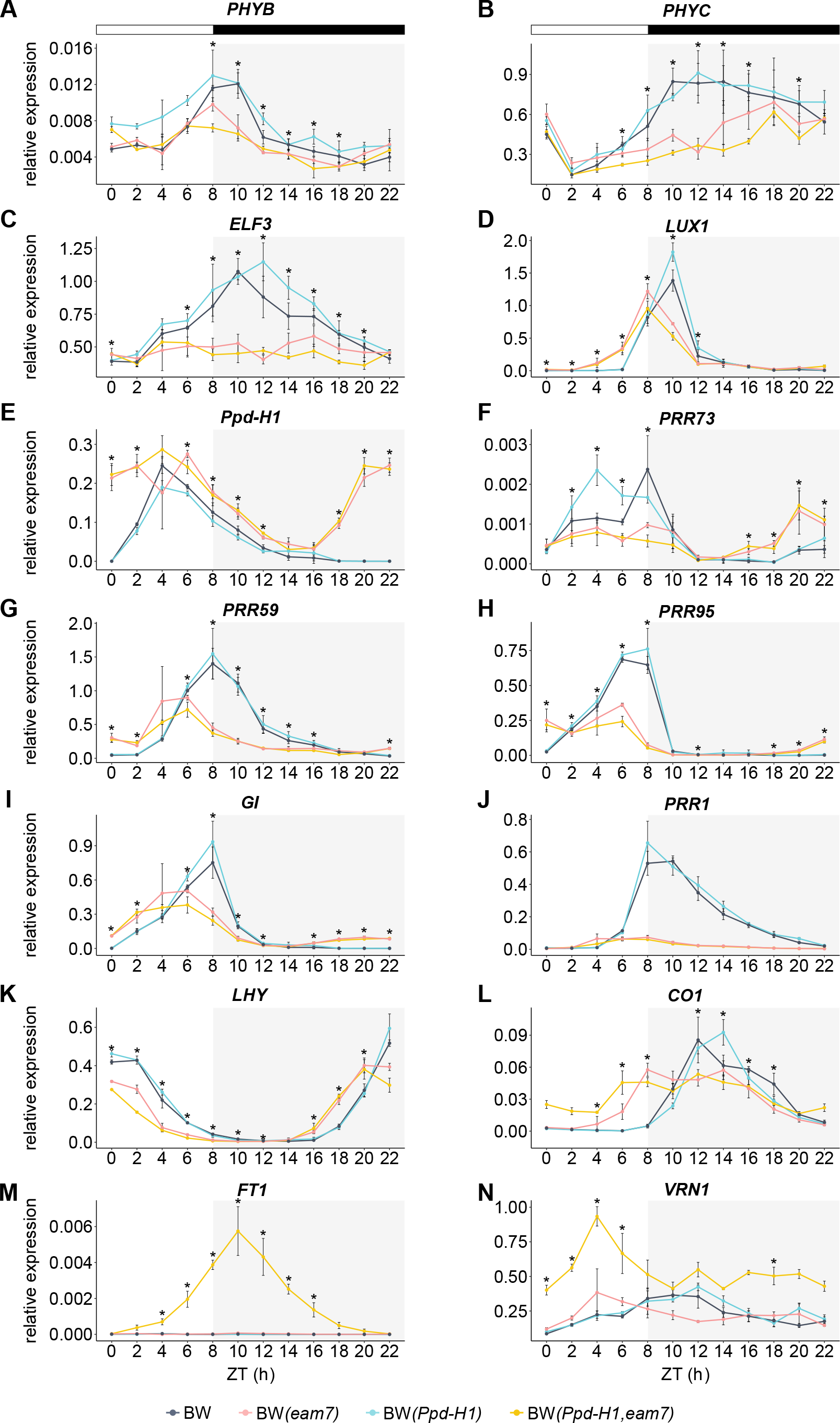
Gene expression pattern of circadian clock genes under SD conditions. Relative expression of *PHYB, PHYC, ELF3, LUX1, Ppd-H1, PRR73, PRR59, PRR95, GI, PRR1, LHY, CO1, FT1* and *VRN1* in Bowman (grey), BW*(Ppd-H1)* (blue), BW*(eam7)* (pink), and BW*(Ppd- H1,eam7)* (yellow) under short-day conditions. Plants were sampled every 2 h over 24 h. White bars indicate day and black bars indicate night. Each value represents the mean of three independent biological replicates, each consisting of two pooled plants. Error bars indicate the standard deviation of the mean; significant differences are indicated by asterisks (* *p* ≤ 0.05) comparing BW and BW*(Ppd-H1)* to BW*(eam7)* and BW*(Ppd-H1,eam7)* **(A-L, N)** or BW, BW*(Ppd-H1)* and BW*(eam7)* to BW*(Ppd-H1,eam7)* **(M)** with Student’s *t*-test, n = 3

Since *eam7* caused the diurnal misregulation of circadian genes and the central photoperiod response gene *Ppd-H1*, we further tested the expression of central floral activators in the photoperiod response pathway of barley (Campoli et al., 2012). *CO1* expression occurred only during the dark period in BW and BW*(Ppd-H1)* but was advanced by 4-6 h and peaked at the end of the light period in BW*(eam7)* and BW*(Ppd-H1,eam7)* (Fig. 3L). *CO2* transcripts could not be detected at any time point during the day in any of the genotypes. *FT1,* typically only expressed under LDs, showed an expression peak under SD only in BW*(Ppd-H1,eam7),* while no transcripts could be detected in the other three genotypes (Fig. 3M). *FT1* expression under SD was thus controlled by *eam7* together with allelic variation at *Ppd-H1*. The upregulation of *FT1* in BW*(Ppd-H1,eam7)* correlated with significantly higher transcript levels of the *MADS*-box gene and floral inducer *VRN1* compared to BW, BW*(eam7)*, and BW*(Ppd-H1)* (Fig. 3N).

Because *FT1* was only expressed in BW*(Ppd-H1,eam7),* but also BW*(eam7)* flowered earlier under SD, we tested the expression of *FT1* and its close homologs *FT2* and *FT3* at later time points during development. We sampled leaf material every 1-2 weeks between 20 and 60 DAE at ZT9 (1 h after dusk) in BW and the introgression lines and tested *FT1*, *FT2*, *FT3,* and *VRN1* transcript levels. However, we could not detect any *FT1* transcripts during development in BW*(eam7)*. We could also not link differences in expression levels in any of the additionally tested genes to the early flowering of BW*(eam7)* under SD compared to BW (Supplemental Fig. S3A-D).

In conclusion, clock genes and phytochromes displayed marked alterations in diurnal expression patterns. This suggests that the gene underlying *eam7* is either a component of the circadian clock or is involved in the light-driven entrainment of the barley circadian clock. The *eam7*-controlled misexpression of clock genes, together with functional variation in *Ppd-H1,* were linked to differences in the diurnal expression of *Ppd-H1* and *CO1,* and the upregulation of *FT1* in BW*(Ppd-H1,eam7)* under SD conditions.

### Biparental mapping identifies *LIGHT-REGULATED WD 1 (LWD1)* as a candidate gene for *eam7*

The *eam7* mutation was mapped to the centromeric region of chromosome 6HS (Stracke and Börner, 1998). For the identification and characterization of *eam7*, the mutant locus *eam7.g* was backcrossed several times to Bowman to generate the introgression lines BW287 (BC_2_) and BW288 (BC_3_, BW*(eam7)*) (Druka et al. (2011). Both lines are early flowering under SD and were thus proposed to carry the same *eam7.g* mutation (Franckowiak and Lundqvist, 2015). Genotyping of BW*(eam7)* (BW288) with the 1536 SNP array identified an introgression of 151.1 cM (BOPA 2_0886 - BOPA 1_1261) on chromosome 6H as the likely location of the causative *eam7* mutation (Druka et al., 2011). We genotyped BW*(eam7)* with the 50k SNP array to confirm the large introgression on 6H, which could be separated into two individual introgressions (Fig. 4A, Supplemental Table S2). The first introgression (6H-1) is spanning 54.61 cM / 380.4 Mbp (position 2.2-378.54 Mbp, Morex V3, Mascher et al., 2021) and contains 812 polymorphic SNPs compared to the recipient parent BW and 2326 high-confidence (HC) genes. The second introgression could be excluded as a likely location of the *eam7* locus since it is located on the long arm of chromosome 6 (Supplemental Table S2).

**Figure 4:**
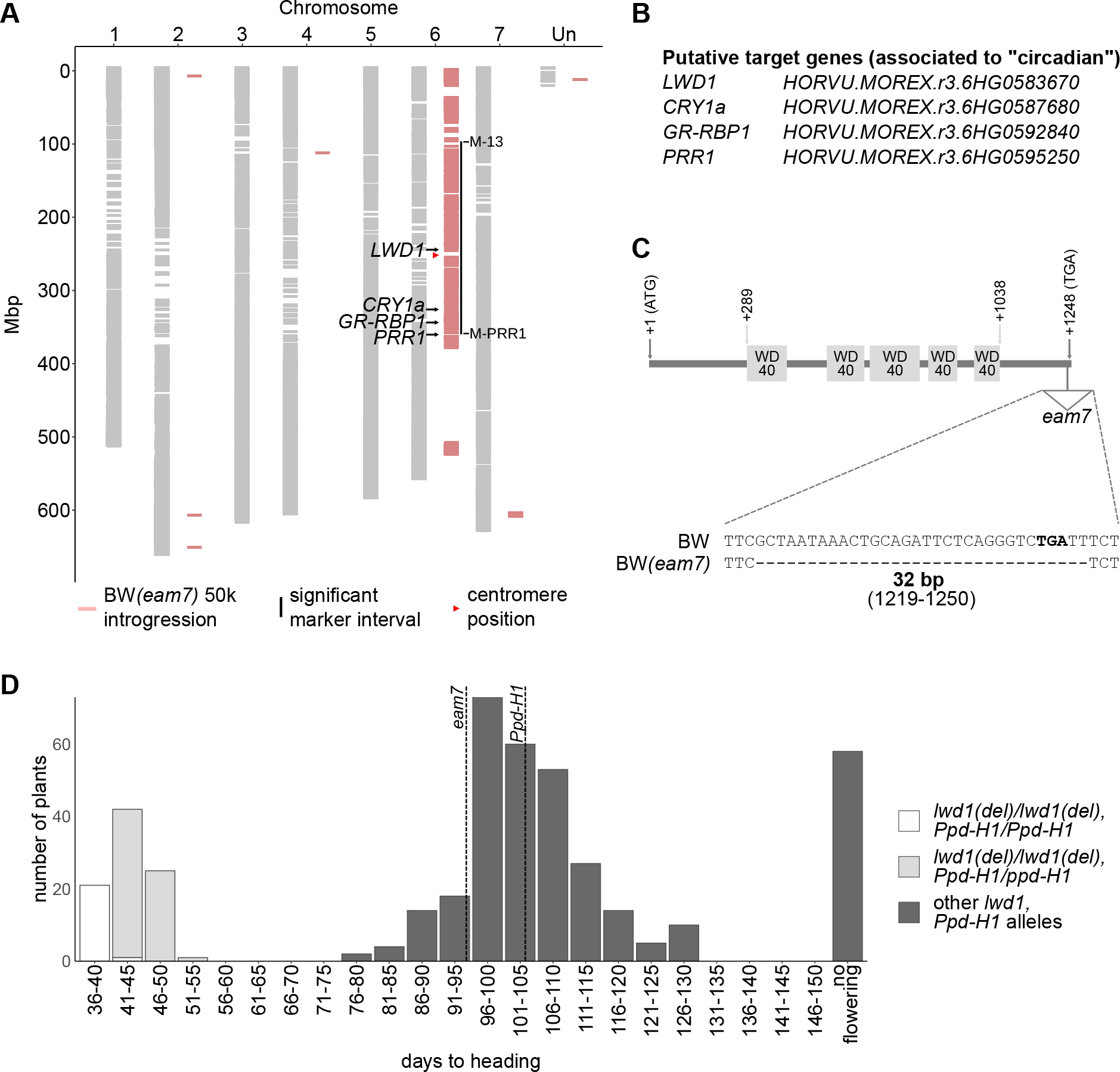
Mapping of *eam7* and identification of *LWD1* as a candidate gene. A. Overview of the *eam7* introgression in parent Bowman, based on 50k Illumina Infinium iSelect SNP array. SNPs were mapped to Morex V3. Horizontal grey bars indicate SNPs that do not differ in Bowman and BW*(eam7),* while pink bars show SNPs polymorphic in BW*(eam7)* compared to Bowman. The vertical black line represents the significant marker interval determined by biparental mapping and its flanking markers (*M-13* and *M-PRR1*), and the red arrow indicates the predicted centromere position (256 Mbp, Mascher et al., 2021). Black arrows show the approximate position of candidate genes. **B** Overview of putative *eam7* candidates related to the term “circadian” within the significant marker interval. **C** Schematic overview of the *LWD1* gene. *LWD1* is a single-exon gene with five W40 repeat domains. The *eam7* allele has a 32 bp deletion at the end of the coding sequence from position 1219 to 1250 (relative to the start codon). The stop codon is indicated in bold. **D** Flowering time under short days of F2 plants of a biparental mapping population, segregating for *eam7* and *Ppd-H1.* This shows 100% co- segregation of the 32 bp *lwd1* deletion *(lwd1 (del))* with early flowering in the presence of a homozygous (white) or heterozygous (light grey) wild-type *Ppd-H1* allele. Plants with other allele combinations (homozygous for the mutated *ppd-H1* allele, heterozygous for *lwd1(del),* or no deletion in *lwd1*) were grouped together (dark grey). Dotted vertical lines indicate the approximate mean flowering time of parents BW*(eam7)* (95.56 ± 5.7 days) and BW*(Ppd-H1)* (106.11 ± 9.7 days). 57 of 423 plants did not flower until the end of the experiment (130 DAE).

To confirm and narrow down the *eam7* location on 6H, we scored 423 F2 plants from a cross between BW*(eam7)* and BW*(Ppd-H1)* for flowering time under SD and genotyped the population for selected SNPs in the introgressed region on 6H (Supplemental Tables S3 and S4). We could confirm that all F2 plants early flowering under SDs carried an introgression on 6H and the wild-type *Ppd-H1* allele on 2H (Supplemental Dataset S1). Further, we could reduce the area of introgression 6H-1 to 2.84 cM / 286 Mbp (position 96.64-382.6 Mbp) containing 1082 HC genes (introgression 6H-1-reduced, Fig. 4A, Supplemental Table S2). These included ten genes homologous to Arabidopsis genes with functions in the circadian clock (Araport 11 annotation, Cheng et al., 2017) (Supplemental Table S5, Supplemental Dataset S2). These candidates included four genes which are all associated with early flowering in barley or Arabidopsis: the clock gene *PRR1 (HORVU.MOREX.r3.6HG0595250),* blue light receptor *CRYPTOCHROME 1a (CRY1a, HORVU.MOREX.r3.6HG0587680), GLYCIN-RICH RNA-BINDING-PROTEIN 1 (GR-RBP1, HORVU.MOREX.r3.6HG0592840)* and *HORVU.MOREX.r3.6HG0583670* with high protein sequence identity (79 and 76%) to Arabidopsis *LIGHT-REGULATED WD 1 (LWD1*) and *LWD2* (Figs. 4B and S4). Due to the higher sequence similarity to *LWD1*, we termed the gene *HvLWD1*.

We sequenced these candidate genes in BW and the derived introgression lines BW*(Ppd-H1)*, BW*(eam7),* and BW*(Ppd-H1,eam7),* the original *eam7*.g mutant Atsel and its parent, Atlas and the derived introgression donor GSHO 579 which was introgressed into Bowman carrying the *eam7* mutation (Druka et al., 2011). *CRY1a* and *GR-RBP1* did not show any SNPs within the coding sequence between BW and BW*(eam7)* and were therefore excluded as candidates for *eam7*. For *PRR1*, we detected two non-synonymous SNPs (T215A and S434P) between BW*(eam7)* and BW (Supplemental Table S5). These SNPs were also present in *eam7* genotypes GSHO 579 and Atsel, but also the parent Atlas. Genotyping *PRR1* in the F2 population revealed one recombinant plant, which was late flowering but carried the SNP haplotype from the introgressed segment (S434P, marker M-PRR1, Supplemental Table S3). We, therefore, excluded *PRR1* as a candidate for *eam7*. For *LWD1*, we identified a 32 bp deletion (position 1219-1252) in BW*(eam7)* and BW*(Ppd-H1,eam7)* but not in BW and BW*(Ppd- H1)* (Fig. 4C, Supplemental Table S5). This deletion is not within a conserved domain but shortens protein length from 415 to 410 aa and was also present in GSHO 579 and the original *eam7* mutant Atsel (Supplemental Fig. S5). The complete co-segregation of *LWD1* with the early flowering phenotype was confirmed by genotyping the F2 population for the 32 bp deletion in *LWD1*. (Fig. 4D, Supplemental Table S3). The deletion, however, was not present in the *eam7* introgression line BW287, suggesting that the causative mutations for early flowering under SDs differed between BW*(eam7)* and BW287.

In summary, the genetic mapping reduced the introgression on chromosome 6H and revealed *LWD1* as a putative candidate gene for *eam7*.

### CRISPR-generated mutants confirm *LWD1* as the gene underlying the *eam7* locus

To confirm *LWD1* as the gene underlying the *eam7* locus, we generated *lwd1* mutant plants using CRISPR-Cas9. GP-fast (spring cultivar Golden Promise with a dominant *Ppd-H1* introgressed from Igri, Gol et al., 2021) was transformed with two different constructs. These targeted either the start or the end of the CDS of *LWD1* to create either a complete knock-out or a mutation similar to the 32 bp deletion present in BW*(eam7)*. From 16 individual mutation events, three homozygous M2 lines were chosen for further experiments: *lwd1-26* and *lwd1- 390,* with mutations early in the coding sequence and a total protein length of 26 and 390 aa, respectively, and *lwd1-402,* with modifications close to the C terminus and a total protein length of 402 aa (Supplemental Fig. S6).

Homozygous M2 and GP-fast plants were grown under SD conditions to test whether the mutant plants were early flowering under SD as observed for *eam7*. All mutant plants flowered early in comparison to GP-fast: *lwd1-26* flowered 45 DAE, followed by *lwd1-402* with 51 DAE and *lwd1-390* with 67 DAE as compared to GP-fast, which flowered 104 DAE (3 plants) or had not flowered (20 plants) when the experiment was terminated (Fig. 5A). *lwd1* shoots and spikes were shorter than in GP-fast, similar to the differences in plant height and spike length between BW*(Ppd-H1,eam7)* and BW (Fig. 5, B and C). While *lwd1* plants produced fewer florets than GP-fast, grain set, and spike fertility significantly increased (Supplemental Fig. S7).

**Figure 5:**
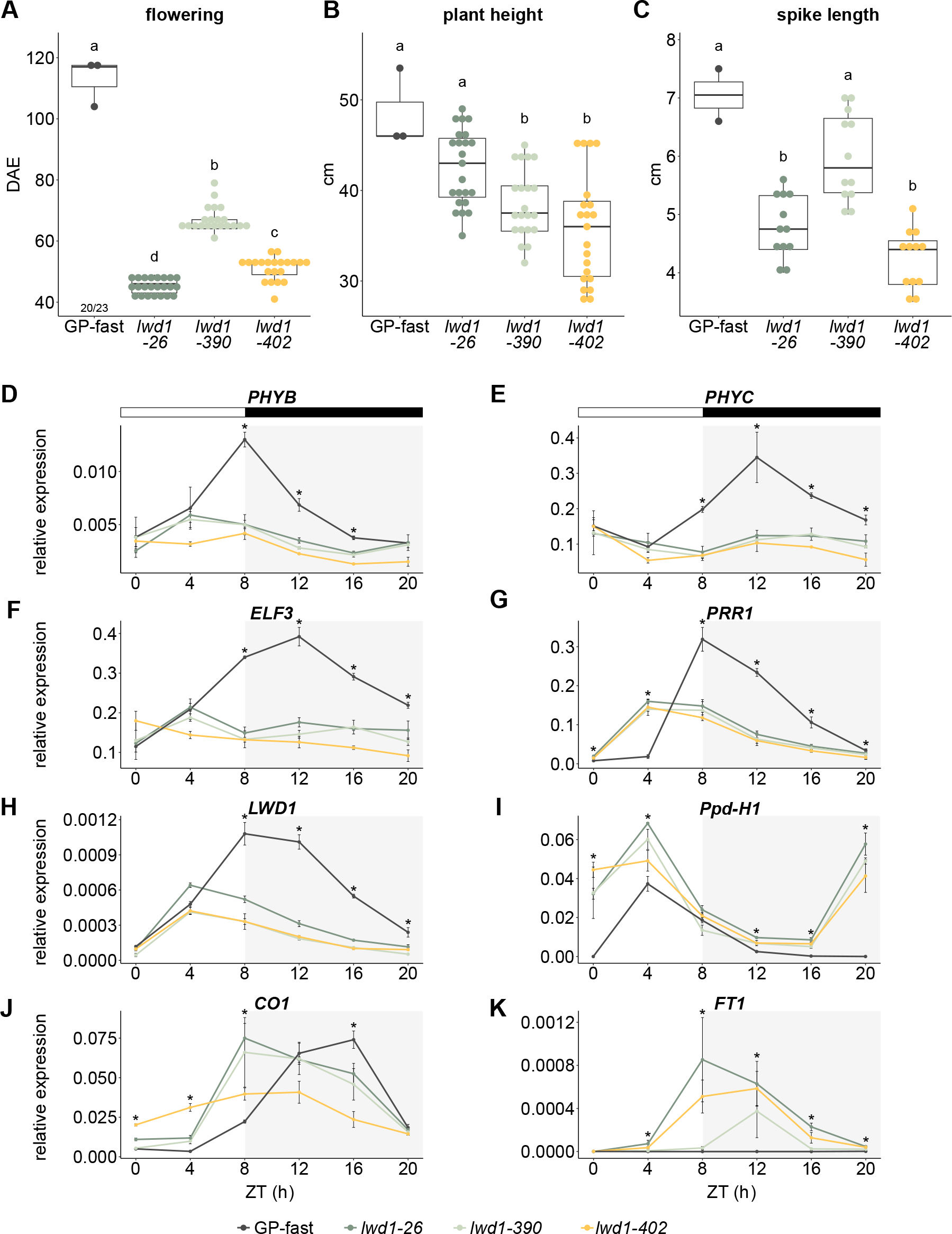
***lwd1* plants phenocopy BW*(Ppd-H1,eam7)*. A-C** Spring cultivar GP-fast and the three *lwd1* mutants *lwd1-26, lwd1-390*, and *lwd1-402* were grown under short-day conditions. Flowering time **(A)**, plant height at flowering (in cm) **(B)**, and spike length (in cm) **(C)** were scored on the main culm. Numbers below boxplots in (A) indicate the number of plants that did not flower until the end of the experiment (130 days after emergence, DAE) compared to the total number of plants scored. Significance levels were determined by one-way ANOVA and subsequent Tukey’s test, *p* ≤ 0.05, n = 3 for GP-fast, n = 12-23 for *lwd1* plants. **D-K** Diurnal gene expression in *lwd1* mutants and wild-type plants. Relative expression of *PHYB, PHYC, ELF3, PRR1, LWD1, Ppd-H1, CO1,* and *FT1* in GP-fast (grey), *lwd1-26* (dark green), *lwd1-390* (light green), and *lwd1-402* (yellow) in short-day conditions. Plants were sampled every 4 h over 24 h. White bars indicate day and black bars indicate night. Each value represents the mean of three independent biological replicates, each consisting of two pooled plants. Error bars indicate the standard deviation of the mean; significant differences are indicated by asterisks (* *p* ≤ 0.05) comparing the three *lwd1* lines to GP-fast with Student’s *t*-test, n = 3

We then tested if the *lwd1* mutant lines were also altered in phytochrome and clock gene expression as observed for the *eam7* genotypes. We grew GP-fast and *lwd1* mutant plants under SD for two weeks and sampled them in 4 h intervals for a complete light-dark cycle. We could confirm that *lwd1* mutant plants were characterized by altered diurnal transcript patterns for phytochromes, clock, and flowering time genes comparable to those observed in *eam7* plants. The expression of phytochromes and *ELF3* showed a strong downregulation in the three *lwd1* mutant lines compared to GP-fast, as seen in *eam7* genotypes compared to BW (Fig. 5D- F). The peak expression of *PRR1* and *LWD1* itself were reduced and advanced by four hours in the *lwd1* mutants compared to GP-fast, similar to *eam7* plants (Figs. 5G, H and S3E). *Ppd-H1* transcription was de-repressed in *lwd1* mutants at night, which increased *Ppd-H1* expression levels in the night and morning (Fig. 5I). In *lwd1 mutants, CO1* expression peaked earlier and at the end of the light phase (Fig. 5J). All *lwd1* lines showed low *FT1* expression levels, whereas no *FT1* transcripts could be detected for GP-fast (Fig. 5K).

To confirm that *lwd1* and *eam7* are allelic, complementation experiments were performed by crossing *lwd1-26*, *lwd1-390,* and *lwd1-402* with BW*(Ppd-H1,eam7)*. The resulting F1 offspring was early flowering under SD, similar to BW*(Ppd-H1,eam7)* plants (Supplemental Fig. S8). We thus concluded that *lwd1* and *eam7* are allelic.

In summary, *lwd1* mutants phenocopied Bowman *eam7* plants. They were early flowering under SD, and the diurnal expression pattern of phytochromes, circadian clock, and flowering genes was altered.

### Natural variation of *LWD1*

*LWD1* belongs to the WD40-repeat proteins, which are conserved across eukaryotes and are involved in highly diverse processes, including flowering and floral development (van Nocker and Ludwig, 2003). In Arabidopsis, two functionally redundant *LWD* proteins, *LWD1* and *LWD2,* control photoperiod response and flowering time under LD and SD (Wu et al., 2008). Barley and other *Triticeae* carry two paralogous genes, *WD40-1,* homologous to *LWD1 and LWD2*, and *WD40-2 (HORVU.MOREX.r3.6HG0604400),* homologous to Arabidopsis *TRANSPARENT TESTA GLABRA 1 (TTG1)* (Strygina and Khlestkina, 2019). Comparison of LWD1 protein sequences across barley, bread and emmer wheat, *Brachypodium distachyon,* rice, sorghum, and maize demonstrated that the amino acid sequences of LWD1 are highly conserved in grasses (Supplemental Fig. S9, A and B). Furthermore, alignment of the last 15 amino acids from HvLWD1 with 281 homologous sequences from plant WD proteins showed that the protein terminus deleted in *eam7* is highly conserved even though located outside the WD repeats (Supplemental Fig. S9C).

We examined natural variation in the coding sequence of *LWD1* by exploiting publicly available exome resequencing data from extensive collections of diverse barley germplasm (Russell et al., 2016; Bustos-Korts et al., 2019). Among the 670 investigated barley accessions, we identified nine haplotypes in a haplotype network analysis (Supplemental Fig. S10, Supplemental Dataset S3). The two major haplotypes (I and II) comprised 97.8% of the analyzed accessions and were the only ones identified in elite barley cultivars. Haplotype I included *LWD1* from the reference cultivar Morex and haplotype II carried a synonymous SNP. Seven additional minor haplotypes (III-IX) were identified in wild and landrace germplasm. Of these, only haplotypes VIII and IX carried non-synonymous changes in *LWD1*. Their effect on protein function was assessed using the Sorting Intolerant from Tolerant (SIFT) algorithm (Sim et al., 2012), which predicted tolerated effects on the protein function for both amino acid substitutions (Supplemental Table S6).

In summary, natural variation in *LWD1* was low, and only a few landraces and wild barley accessions showed variation in *LWD1* with no or predicted low-effect changes in the LWD1 protein. This result suggested that *LWD1* is functionally conserved and under strong selection.

## Discussion

### *LWD1* is a candidate for the *eam7* locus

We identified *LIGHT-REGULATED WD 1 (LWD1)* as a candidate gene underlying the *eam7* locus. We demonstrate that *eam7* in the background of a wild-type *Ppd-H1* allele causes rapid flowering and near day-length neutrality under SDs.

Stracke and Börner (1998) mapped *eam7* close to the centromere on the short arm of chromosome 6H, and Druka et al. (2011) identified an introgression of 151.1 cM on 6H in the background of Bowman as the likely location of the causative *eam7* mutation. We narrowed down the genomic location of *eam7* to 2.84 cM using a biparental mapping population of a cross between the *eam7* introgression line BW*(eam7)* and the wild-type *Ppd-H1* introgression line BW*(Ppd-H1).* Within this genomic region, we identified a 32 bp deletion in the coding sequence of *LWD1* as a candidate polymorphism for the early flowering phenotype in the population (Fig. 4). The deletion is located at the C-terminus of the gene and reduces protein length from 415 to 410 aa. To confirm *LWD1* as a candidate for *eam7*, we used CRISPR-Cas9 to generate different *lwd1* mutants in the genotype GP-fast carrying a wild-type *Ppd-H1* allele. All analyzed *lwd1* mutants flowered significantly earlier than the wild-type under SD conditions, irrespective of the size of the protein truncation (Fig. 5), confirming that *LWD1* controls photoperiodic flowering in barley. The strong effects of C-terminal protein truncations in *eam7* and *lwd1-402*, and the high degree of conservation of the C-terminal amino acids in WD proteins across different taxa, indicate that the terminal part of the LWD1 protein is crucial for its function. In addition, all three *lwd1* mutants displayed the same changes in diurnal expression patterns of core clock genes and clock output genes as observed for *eam7* compared to the wild-type genotypes. Finally, allelic mutants between *eam7* and *lwd1* mutants were early flowering under SD, thus confirming *LWD1* as the gene underlying *EAM7*.

### *EAM7* is important for photoperiod sensing in barley

We demonstrated that *eam7* in the background of a wild-type *Ppd-H1* allele causes rapid flowering and near day-length neutrality under SDs. We thus concluded that *eam7* is important for photoperiod perception in barley. This effect is reminiscent of the *early maturity* mutants *eam5*, *eam8,* and *eam10,* which are also early flowering under non-inductive photoperiods and were identified as mutant alleles of *PHYC* and the core clock components *ELF3* and *LUX1*, respectively (Faure et al., 2012; Campoli et al., 2013; Pankin et al., 2014).

In *eam7* plants, allelic variation at the major photoperiod response gene *Ppd-H1* strongly affected flowering time under SD; *eam7* plants with a wild-type *Ppd-H1* allele flowered more than 50 days earlier than those with a mutated *ppd-H1* allele (Fig. 1). Furthermore, *eam7* plants were characterized by a strong upregulation of *Ppd-H1* in the night and the morning, which was linked to the expression of *FT1* and early flowering under SDs (Fig. 3). We thus propose that *eam7* is an upstream transcriptional regulator of *Ppd-H1*. The upregulation of *Ppd-H1* in the night has already been correlated to *FT1* expression under SDs and to day-neutral flowering in *eam5*, *eam8,* and *eam10* and photoperiod insensitive wheat lines (Shaw et al., 2012; Faure et al., 2012; Campoli et al., 2013; Pankin et al., 2014). *eam7* thus likely alters photoperiodic flowering by controlling the diurnal expression pattern of *Ppd-H1*. However, in contrast to *eam8 (elf3),* which causes day-neutral flowering irrespective of allelic variation at *Ppd-H1, eam7* has only a minor effect on flowering time in the background of a mutated *ppd-H1* allele. Early flowering under SDs in *eam7* mutants was thus dependent on the presence of the wild-type *Ppd-H1* allele and the night-time upregulation of *Ppd-H1*.

It has been demonstrated that the repression of *Ppd-H1* at night is controlled by *ELF3 (EAM8)* and *LUX1 (EAM10)* in barley and wheat (Faure et al., 2012; Campoli et al., 2013; Alvarez et al., 2023). In Arabidopsis, *ELF3* and *LUX1* interact to bind the promoter of the *Ppd-H1* ortholog *PRR7* to repress the expression during the night (Mizuno et al., 2014), and this function is likely conserved in barley and wheat (Faure et al., 2012; Campoli et al., 2013; Alvarez et al., 2023). *ELF3* expression was strongly downregulated in *eam7* plants suggesting that *LWD1* controls *Ppd-H1* expression through *ELF3* (Fig. 3). However, *EAM7* might also directly regulate *Ppd-H1* expression, since in Arabidopsis the close paralogs *LWD1* and *LWD2* directly bind to the promoters of *PRR* genes (Wang et al., 2011). In contrast to *EAM7* in barley, Arabidopsis *LWD1* and *LWD2* act as positive regulators of *PRR* expression (Wang et al., 2011). In addition to *ELF3*, we also observed the downregulation of *PHYC* and *PHYB* in *eam7* plants. *PHYB* and *PHYC* are necessary for the light activation of *Ppd-H1* and act as upstream repressors of *ELF3* in barley, wheat, and *Brachypodium* (Chen et al., 2014; Pankin et al., 2014; Alvarez et al., 2023; Woods et al., 2023). *EAM7* might therefore affect flowering time by modifying the expression of phytochromes and *ELF3* and, thus, the light input into the photoperiod pathway.

In *Arabidopsis thaliana*, *LWD1* and *LWD2* regulate photoperiodic flowering by advancing the expression phase of core clock and clock output genes under light/dark conditions (Wu et al., 2008). The early flowering phenotype of the Arabidopsis *lwd1lwd2* double mutant was attributed to a phase shift of the clock target and central photoperiod response gene *CONSTANS* (*CO*) and a consequent increase in *FT* expression. Similarly, in *eam7* mutants, the expression phase of evening expressed clock genes *PRR59/95* and *GI* shifted approximately four hours forward, suggesting that *LWD1* controls the expression phase of the central oscillator genes in barley (Fig. 3). This phase shift was correlated to a forward shift and day-time expression of *CO1* under SDs in the *eam7* mutants (Fig. 3). In Arabidopsis, the coincidence of *CO* expression with the light period is crucial for stabilizing the protein and expressing the florigen *FT* (Sawa et al., 2007; Jang et al., 2008). Similar to the Arabidopsis *lwd1lwd2* double mutants, the *eam7* mutants with an altered diurnal expression of *CO1* were characterized by *FT1* expression in non-inductive photoperiods. However, in barley and wheat, *CO1* mainly acts as a weak heading time repressor and accelerates flowering only in the absence of *Ppd-H1* (Shaw et al., 2020). Furthermore, night-break experiments have revealed that the length of the night and not of the light period is critical for the perception of inductive photoperiods in monocots (Pearce et al., 2017; Gao et al., 2019). Furthermore, BW*(eam7)* also displayed a shift in *CO1* phase expression into the day as observed in BW*(Ppd-H1,eam7),* but no *FT1* expression and only a minor acceleration in flowering under SDs. Nevertheless, the advance in phase expression of clock and clock output genes was comparable between barley *eam7* mutants and Arabidopsis *lwd1lwd2* double mutants, and the function of *LWD1* in controlling the light entrainment of the clock is thus likely conserved across these taxa.

In summary, we have successfully identified *LWD1* as a promising candidate to underly the *eam7* locus. We propose that *LWD1* functions as an upstream activator of the night-time repressor *ELF3* in the light entrainment pathway of the barley circadian clock. *eam7* plants were early flowering in non-inductive photoperiods due to the reduced activation of *ELF3*, the upregulation of *Ppd-H1* during the night, and consecutive *FT1* expression under SDs. *LWD1* is an interesting target to modulate photoperiod sensitivity to breed for barley cultivars adapted to short growing seasons.

## Methods

### Plant material

Spring cultivar Bowman (BW) with a mutated *ppd-H1* allele and three Bowman-derived introgression lines were used in this study. The introgression line BW281 (GSHO 1872) carries a wild-type *Ppd-H1* allele introgressed from winter barley KT1031 (GSHO 1568) (Druka et al., 2011). BW288 (GSHO 2068, NGB 20572) is an introgression line with the *eam7.g* mutation introgressed from Club Mariout/6*California Mariout (GSHO579, Gallagher et al., 1991) in Bowman by three rounds of backcrossing (Druka et al., 2011). We termed these introgression lines BW*(Ppd-H1)* and BW*(eam7)*. We crossed BW*(Ppd-H1)* and BW*(eam7)* to generate a line with a wild-type *Ppd-H1* allele and the mutation at *eam7.g* and termed this cross BW*(Ppd- H1,eam7)*.

The F2 generation of the cross between BW*(Ppd-H1)* and BW*(eam7)* was used for segregation analysis and mapping of *eam7*. Furthermore, the *eam7* introgression donor GSHO579 and the original *eam7.g* mutant Atsel (CIho 6250) and its genetic background Atlas (PI 539108) were included to sequence potential candidate genes. We also included BW287 (NGB 20571), which was generated by backcrossing Club Mariout/6*California Mariout with *eam7.g* into Bowman by two cycles of crossing and was thus reported to be allelic to BW*(eam7)* (Druka et al., 2011).

The introgression line GP-fast, spring cultivar Golden Promise with a dominant *Ppd-H1* allele introgressed from winter cultivar Igri (Gol et al., 2021), was transformed to generate mutants in *LIGHT-REGULATED WD 1 (LWD1, HORVU.MOREX.r3.6HG0583670)* using CRISPR-Cas9.

Three independent homozygous M2 lines, *lwd1-26, lwd1-390,* and *lwd1-402,* were selected for functional analyses and crossing to BW*(Ppd-H1,eam7*). *lwd1-26* has two deletions within the CDS (-C21, -C23) and one single insertion (+T114) that lead to a frameshift and premature stop codon, reducing the protein from 415 (WT) to 26 amino acids (aa). *lwd1-390* has two deletions (66 bp, position 15-80 and 9 bp, position 109-117) that are in frame and lead to the deletion of amino acids 6-27 and 37-39, reducing the protein size to 390 aa. *lwd1-402* has a 39 bp deletion (position 1198- 1238) that reduces protein length to 402 aa.

### Plant phenotyping and growth conditions

BW and the derived introgression lines BW*(Ppd-H1)*, BW*(eam7),* and BW*(Ppd-H1,eam7)* were cultivated under controlled growth conditions for phenotyping and gene expression analyses. All plants were grown in soil (Einheitserde ED73, Einheitserde Werkverband e.V., with 7% sand and 4 g/L Osmocote Exact Hi.End 3-4M, 4^th^ generation, ICL Group Ltd.) and were stratified for four days in 4°C before moving them to controlled growth conditions.

For phenotyping, plants were grown in plant growth chambers under long-day (LD, 16 h light, 20°C, PAR ∼250 µmol/m^2^s; 8 h dark, 16°C) and short-day (SD, 8 h light, 20°C, PAR ∼250 µmol/m^2^s; 16 h dark, 16°C) conditions in QuickPot E 24/10 trays (HerkuPlast Kubern GmbH). Flowering was scored in days after emergence (DAE) as the period between emergence from soil and reaching Zadoks stage 49 when the awns exited the leaf sheath (Zadoks et al., 1974). Plant height, leaf number on the first emerging shoot (main culm), and tiller number were scored at flowering. Plant height was scored as the distance from soil to the flag leaf ligule of the main culm. Leaf number was scored on the main culm, and tiller number as all tillers emerging after the main culm. The length and width of the flag leaf on the main culm were measured as the leaf blade length (from the ligule to the leaf tip) and the maximum width of the blade under SDs. Spike length, floret number, and grain number of the main culm were scored at maturity. Fertility was calculated as the percentage of florets on the mature spike that developed into grains. The experiment was stopped 125 DAE, and all plants that had not flowered up to this point were scored as “not flowering”.

Main shoot apex (MSA) development of all four genotypes was monitored and quantified based on the scale by Waddington et al. (1983). Once or twice a week, the development of the MSA of the main stem of four randomly chosen plants per genotype was dissected. MSA development was documented using the stereo microscope Nikon SMZ18 with a Nikon DS-Fi2 camera and was analyzed with the NIS-Elements Software (version 5.21.03, Nikon Instruments Europe BV). Under LD, plants were dissected every 2 to 10 days starting from 5 DAE until pollination (W10.0). Under SD, plants were dissected every 6 to 8 days between 6 and 47 DAE and every 4 to 13 days from 48 DAE until pollination. The number of developing spikelet meristems (SM), including those that had initiated floret meristems (FM) or developed into florets, was determined on the main inflorescence. The R package *segmented* (version 1.6-2, Muggeo, 2008) was used to calculate broken-line regressions for SM initiation and FM abortion by plotting the number of SM against the Waddington stage. One breakpoint was calculated automatically to separate initiation from abortion and was set as the maximum SM number stage. The corresponding maximum SM number was calculated using linear regression calculated with *segmented*.

The *lwd1* mutant and wild-type plants were grown in controlled plant growth chambers under SD in QuickPot 96T trays (HerkuPlast Kubern GmbH) until maturity. Flowering, plant height, leaf and tiller number, leaf size, and yield parameter were scored as described above. Plants that did not flower until 130 DAE were scored as “not flowering”.

### Gene expression analysis

Gene expression analysis was performed in two independent experiments in Bowman and derived introgression lines BW*(Ppd-H1),* BW*(eam7),* and BW*(Ppd-H1,eam7)* and in the CRISPR-Cas9-generated mutants *lwd1-26, lwd1-390, lwd1-402* and wild-type GP-fast.

Plants were sown in QuickPot 96T trays (HerkuPlast Kubern GmbH) and transferred to plant growth chambers after four days of stratification at 4°C. Plants were cultivated under SD conditions until 14 DAE. BW and derived introgression lines were sampled every two hours, the *lwd1* mutants and wild-type GP-fast every four hours over a complete light/dark cycle of 24 hours (light from ZT0-8, dark from ZT8-24). For each replicate, the middle sections of the youngest, fully elongated leaf of two plants were pooled. Three biological replicates were sampled, frozen immediately in liquid nitrogen, and stored at -80°C until further analysis. In addition, the middle sections of the youngest, fully elongated leaf of the same genotypes were sampled at ZT9 once a week for six weeks, starting at 19 DAE. These leaf samples were used to monitor *FT1*, *FT2*, *FT3*, and *VRN1* transcript levels during plant development.

RNA was extracted by grinding the samples using TRIzol (Thermo Fisher Scientific) according to the manufacturer’s instructions. RNA was resuspended in 60 µl of DEPC-treated water at 4°C overnight. Remaining DNA was removed by subsequent DNAse I treatment (Thermo Fisher Scientific) according to the manufacturer’s instructions. cDNA was synthesized on 2 µg of total RNA using ProtoScript II First Strand cDNA Synthesis Kit (NEB) following the manufacturer’s instructions. Gene expression levels were determined by qRT-PCR in a LightCycler 480 (Roche) using gene-specific primers (Supplemental Table S7). The reaction was performed using 4 µl of cDNA, 5 µl of 2X Luna qPCR Master Mix (NEB), 0.02 mM of forward and reverse primer, and 0.75 µl of water with the amplification conditions 95°C for 5 min, 40 cycles of 95°C (10 s), 60°C (10 s) and 72°C (10 s). Non-template controls were added to each plate, and dissociation analysis was performed at the end of each run to confirm the specificity of the reaction. Starting amounts for transcript levels were calculated based on the titration curve for each target gene using the LightCycler 480 Software (Roche; version 1.5.1.62). Two technical replicates were used and averaged in analyses for each biological replicate. The expression of *Actin* was used as a reference to calculate the relative gene expression of the target genes.

### Identification of a candidate gene underlying *eam7*

DNA was extracted from BW*(eam7)* leaf using the DNeasy Plant Mini Kit (QIAGEN) according to the manufacturer’s instructions, and plants were genotyped with the 50k Illumina Infinium iSelect SNP Array (Bayer et al., 2017). The *eam7* introgression area was visualized by comparing the BW*(eam7)* allele to the Bowman allele for each SNP. SNPs were plotted against their respective position within the Morex V3 genome assembly in Megabase pairs (Mbp, Mascher et al., 2021). Ambiguous (including heterozygous) and failed SNPs were removed.

For segregation analyses, the Bowman introgression lines BW*(eam7)* and BW*(Ppd-H1)* were crossed, and 423 F2 plants together with nine plants each of BW, BW*(eam7),* and BW*(Ppd-H1)* were sown in QuickPot E24/10 trays. After four days of stratification at 4°C, plants were cultivated under SD conditions, and DNA was extracted using the KingFisher Flex (Thermo Fisher Scientific) and the BioSprint 96 DNA Plant Kit (QIAGEN). Flowering time was scored as described above. Plants that did not flower until the end of the experiment (130 DAE) were scored as “not flowering”.

Based on the introgression area determined with the 50k SNP array, several CAPS markers were designed on chromosome 6 with indCAPS and Primer3Plus to localize *eam7* (Untergasser et al., 2007; Hodgens et al., 2017). In addition, the CAPS marker designed by Turner et al. (2005) was used to determine whether the plants carried a wild-type or mutated *Ppd-H1* allele. Co-segregation of early flowering with a 32 bp deletion in *LWD1* was tested by amplifying the surrounding area with the primer pair *lwd1-del* and comparing fragment sizes on an agarose gel. All used CAPS markers and primers are listed in Supplemental Table S4.

*eam7* candidate genes *CRY1a, GR-RBP1, LWD1,* and *PRR1* were amplified in Bowman and the derived introgression lines BW*(Ppd-H1)*, BW*(eam7),* BW287 and BW*(Ppd-H1,eam7)*, in GSHO 579 and the original *eam7* mutant Atsel and its parent Atlas. The full genomic sequence was Sanger sequenced to identify mutations. Primers used for amplification and Sanger sequencing can be found in Supplemental Table S8.

### Generating *lwd1* mutants using CRISPR-Cas9

To confirm *LWD1* as a candidate gene for *eam7, lwd1* mutants were generated using CRISPR- Cas9. The vector system by Kumar et al. (2018) was used to design transformation vectors targeting *LWD1*. Two approaches were used: Approach 1 included two guide RNAs (gRNAs) targeting the start of the CDS of *LWD1* (CDS position 7 and 97). The second approach targeted the end of the coding sequence (CDS position 1182 and 1222, Supplemental Table S9) to generate mutations comparable to BW*(eam7)*. gRNAs were designed using RGEN Tools Cas-Designer (Bae et al., 2014; Park et al., 2015). Cloning was performed according to the protocol by Kumar et al. (2018): The single gRNA strands were hybridized and cloned into the shuttle vectors pMGE625 or pMGE627 by a BpiI cut/ligation reaction. A second cut/ligation reaction (BsaI) was used to transfer the gRNA transformation units (TUs) to the recipient vector pMGE599. The final vectors were used to transform GP-fast via embryo transformation according to the protocol by Hensel et al. (2009). Successful insertion of the transformation vector into the genome was tested by PCR (primer Hyg-156 and Hyg-047, Supplemental Table S8) on M0 plants. M2 plants were genotyped for mutations by amplifying the entire genomic sequence of *LWD1* and subsequent Sanger sequencing (Supplemental Table S8). Three lines that showed different mutations were selected for further experiments and were termed *lwd1-26, lwd1-390*, and *lwd1-402*.

### Allelism tests

Allelism tests were performed by generating F1 crosses of the mutant lines *lwd1-26, lwd1-390,* and *lwd1-402* with BW*(Ppd-H1, eam7).* As controls, plants were crossed with BW*(Ppd-H1)* and GP-fast. 5-10 plants per cross of the resulting F1 generation were grown in 7x7x7.5 cm pots with the parental plants under SD conditions in plant growth chambers, and flowering time was scored as described above. The experiment was terminated after 125 DAE and all plants that did not flower were scored as “not flowering”. DNA was extracted from all F1 and parent plants using the KingFisher Flex (Thermo Fisher Scientific) and the BioSprint 96 DNA Plant Kit (QIAGEN). The complete *LWD1* CDS was amplified by PCR and sequenced using Sanger sequencing with the same primers described above (Supplemental Table S8).

### Natural variation

A haplotype network analysis was conducted based on combined exome resequencing data from Russell et al. (2016) and the WHEALBI collection (Bustos-Korts et al., 2019). This combined set includes 213 cultivars, 303 landraces, 111 wild barley accessions (*H. vulgare* ssp. *spontaneum*), one *H. agriocrithon* accession, and 42 *H. vulgare* spp. *vulgare* accessions with unassigned breeding history (referred to as ’unknown’). The haplotype network was constructed as described by Walla et al. (2020).

Orthologs of HvLWD1 in other grasses were identified in the Ensembl Plants database (Bolser et al., 2016). The multiple protein sequence alignment was performed using CLUSTAL Omega (1.2.4) (Madeira et al., 2022), and conserved domains were identified with NCBI conserved domains (Marchler-Bauer et al., 2017). Sequence conservation analysis was performed as described in Pankin et al. (2014). The last 15 amino acids from HvLWD1 were used to extract 281 sequences of proteins annotated as WD proteins from plants using NCBI Blastp (e-value cut-off: 0.05). Sequences were mapped using MAFFT v7 (“auto” method, Katoh et al., 2019), and the sequence logo was visualized with WebLogo 3 (Crooks et al., 2004).

### Statistical analyses

All statistical tests were performed using R Studio (RStudio Team, 2022). A two-tailed, unpaired Student’s *t*-test (function *t_test* from the package *rstatix,* v0.7.2) was used to determine the significance between two group means, with a *p*-value cut-off at ≤ 0.05. Significance between more than two groups was determined using a one-way ANOVA (function *aov*) and a subsequent Tukey test (function *HSD.test* from package *agricolae*, v1.3-5), *p*-value cut-off at ≤ 0.05. Polynomial regressions (Loess smooth line) were calculated with a 95% confidence interval.

## Acknowledgements

We would like to thank Einar B. Haraldsson for providing the BLASTp results of barley against *Arabidopsis thaliana* and Kumsal Ecem Çolpan Karışan for the critical reading of the manuscript. The funders had no role in study design, data collection and interpretation, or the decision to submit the work for publication.

## Author contributions

GH and MvK conceived and designed the experiments. GH conducted plant phenotypic analyses, generated BW*(Ppd-H1,eam7)* crosses, designed CAPS markers, cloned CRISPR- Cas9 transformation vectors, did sequence analyses, and analyzed the data. AW performed the haplotype analysis. TR and GH performed qPCR experiments and genotyping of the mapping population. GB and GöH transformed plants and regenerated *lwd1* mutants. RS generated allelic *lwd1* crosses. GH wrote the manuscript with the help of AW and MvK.

